# Gene activation precedes DNA demethylation in response to infection in human dendritic cells

**DOI:** 10.1101/358531

**Authors:** Alain Pacis, Florence Mailhot-Léonard, Ludovic Tailleux, Haley E Randolph, Vania Yotova, Anne Dumaine, Jean-Christophe Grenier, Luis B Barreiro

## Abstract

DNA methylation is considered to be a relatively stable epigenetic mark. Yet, a growing body of evidence indicates that DNA methylation levels can change rapidly, for example, in innate immune cells facing an infectious agent. Nevertheless, the causal relationship between changes in DNA methylation and gene expression during infection remains to be elucidated. Here, we generated time-course data on DNA methylation, gene expression, and chromatin accessibility patterns during infection of human dendritic cells with *Mycobacterium tuberculosis*. We found that the immune response to infection is accompanied by active demethylation of thousands of CpG sites overlapping distal enhancer elements. However, virtually all changes in gene expression in response to infection occur prior to detectable changes in DNA methylation, indicating that the observed losses in methylation are a downstream consequence of transcriptional activation. Footprinting analysis revealed that immune-related transcription factors (TF), such as NF-κB/Rel, are recruited to enhancer elements prior to the observed losses in methylation, suggesting that DNA demethylation is mediated by TF binding to cis-acting elements. Collectively, our results show that DNA demethylation is not required for the establishment of the core regulatory program engaged upon infection.

## INTRODUCTION

Innate immune cells, such as dendritic cells (DCs) and macrophages, are the first mediators recruited in response to an invading pathogen. Upon stimulation, these cells considerably shift their transcriptional program, activating hundreds of genes involved in immune-related processes in a rapid and highly choreographed fashion. This is achieved through the binding of signal-dependent transcription factors (TFs), including NF-κB/Rel, AP-1, and interferon regulatory factors (IRFs), to gene regulatory regions of the genome where recruitment of various co-activators is initiated [1, 2]. Alterations to the epigenome, such as histone modifications and DNA methylation, are recognized as important permissive or suppressive factors that play an integral role in modulating access of TFs to cis-acting DNA regulatory elements via the regulation of chromatin dynamics. Consequently, changes to the epigenetic landscape are expected to have a significant impact on gene expression.

Many studies have highlighted the importance of histone modifications in regulating complex gene expression programs underlying immune responses [3, 4]. However, the exact role that DNA methylation plays in innate immune response regulation remains ambiguous. We have previously shown that infection of post-mitotic DCs is associated with an active loss of methylation at enhancers and that such demethylation events are strongly predictive of changes in expression levels of nearby genes [5]. Many other studies correlate these two processes [6-13], but it remains unclear whether altered methylation patterns directly invoke transcriptional modulation or whether such patterns are the downstream consequence of TF binding to regulatory regions. Thus, the causal relationship between changes in DNA methylation and gene expression during infection remains unresolved. To address this question, we characterized in parallel genome-wide patterns of DNA methylation, gene expression, and chromatin accessibility in non-infected and *Mycobacterium tuberculosis* (MTB)-infected DCs at multiple time points. Our results show that the loss of DNA methylation observed in response to infection is not required for the activation of most enhancer elements and that, instead, demethylation is a downstream consequence of TF binding.

## RESULTS

### Bacterial infection induces stable DNA demethylation at enhancers of dendritic cells

To investigate the relationship between changes in gene expression and DNA methylation in response to infection, we infected monocyte-derived DCs from 4 healthy individuals with a live virulent strain of *Mycobacterium tuberculosis* (MTB) for 2-, 18-, 48-, and 72-hours. At each time-point, we obtained single base-pair resolution DNA methylation levels for over 130,000 CpG sites using a customized capture-based bisulfite sequencing panel (SeqCap Epi, see **Methods**), in matched non-infected and MTB-infected DCs. Our customized SeqCap Epi panel interrogates 33,059 regions highly enriched among putative enhancer elements (58% are associated with the H3K4me1 enhancer mark [14]; **Supplementary Figure 1A**), which are the main targets of methylation changes in response to infection [5]. In total, we generated ~717 million single-end reads (mean = 17.5 million reads per sample; **Supplementary Table 1**), resulting in an average coverage of ~70X per CpG site (**Supplementary Figure 1B**). Methylation values between samples were strongly correlated, attesting to the high quality of the data (**Supplementary Figure 1C**; median r across all samples = 0.94).

We next assessed temporal changes in methylation levels in response to infection using the DSS software [15]. We defined differentially methylated (DM) CpG sites as those showing a significant difference of methylation between infected and non-infected samples at a False Discovery Rate (FDR) < 0.01 and an absolute mean methylation difference above 10%. Using these criteria, we identified 6,174 DM CpG sites across the time course of infection. Consistent with previous findings [5], the vast majority of changes in methylation (87%) were associated with the loss of DNA methylation in infected cells (**Figure 1A,B**).

**Figure 1.**
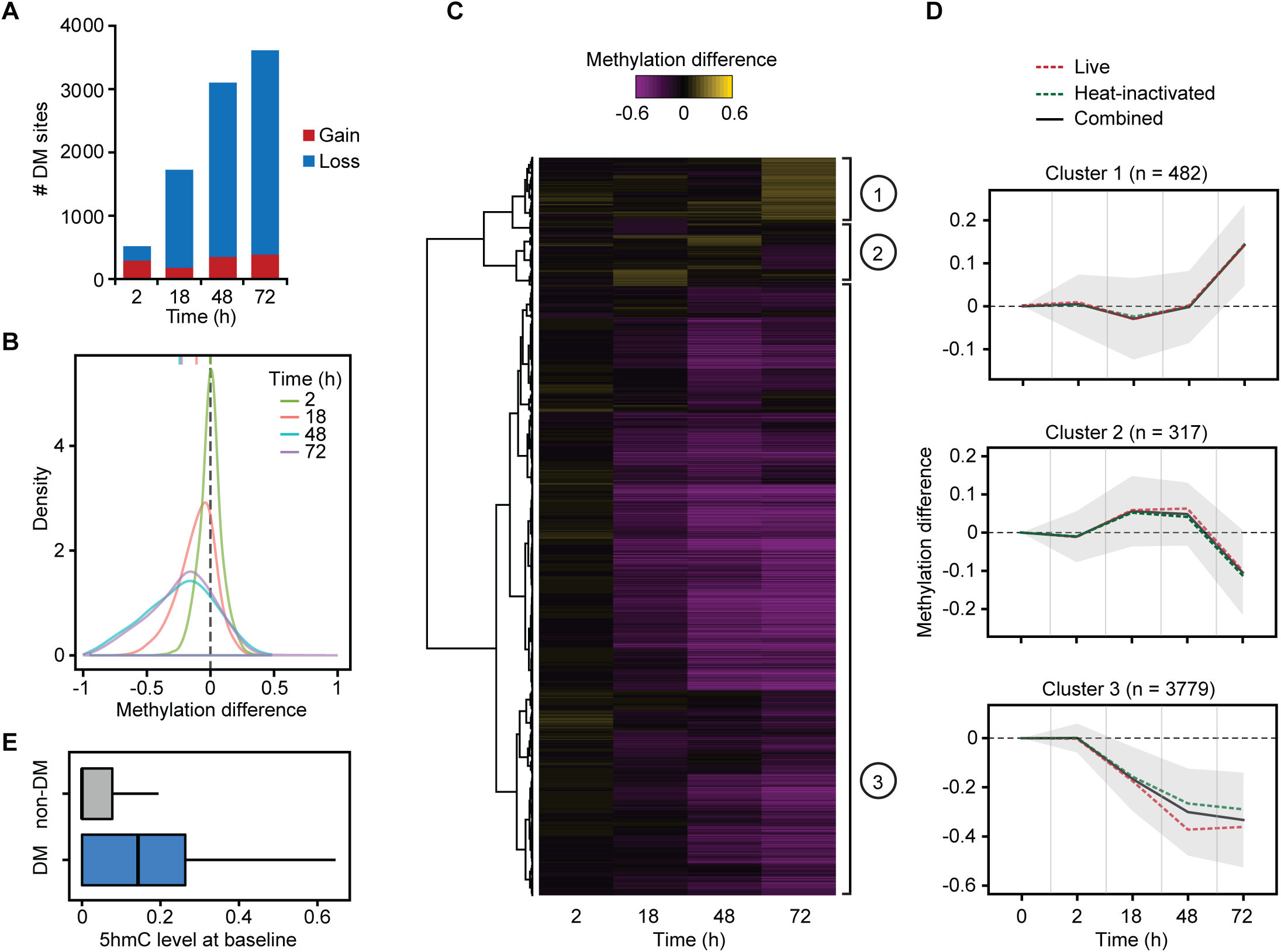
(**A**) Barplots showing the number of differentially methylated (DM) CpG sites identified at a |methylation difference| > 10% and FDR < 0.01 (y-axis) at each time point after MTB infection (2, 18, 48, and 72 hours (h)) (x-axis). (**B**) Distribution of differences in methylation between infected and non-infected cells at DM sites, at each time point. (**C**) Heatmap of differences in methylation constructed using unsupervised hierarchical clustering of the 4,578 DM sites (identified at any time point using live and heat-inactivated MTB-infected samples combined; y-axis) across four time points after infection, which shows three distinct patterns of changes in methylation. (**D**) Mean differences in methylation of CpG sites in each cluster across all time points; shading denotes ±1 standard deviation. For visualization purposes, we also show the ‘0h’ time point, where we expect no changes in methylation. (**E**) Boxplots comparing the distribution of 5hmC levels in non-infected DCs between non-DM and DM sites (Cluster 3).

To test if live bacteria were required to induce the observed changes in DNA methylation, we collected similar data on DCs exposed to heat-killed MTB in addition to the live MTB experiments. Changes in methylation in response to live and heat-killed MTB were strikingly correlated, particularly at later time-points post-infection (r ≥ 0.84 at 18h and above; **Supplementary Figure 2**). These results show that DCs do not require exposure to a live pathogen to elicit the overall demethylation detected in response to infection. Simply, the engagement of innate immune receptors and activation of pathways involved in pathogen sensing and elimination is sufficient to induce methylation shifts. Hierarchical clustering analysis of the DM sites observed when considering samples exposed to either live or heat-killed bacteria showed that >80% of the sites exhibited a gradual loss of methylation over the time course of infection until methylation marks were almost completely erased and that very few changes were detectable at 2 hours post-infection (DM Cluster 3; **Figure 1C,D**; **Supplementary Table 2**).

Monocyte-derived DCs do not proliferate in response to infection [5] and, therefore, any observed losses in methylation must occur through an active mechanism involving the ten-eleven translocation (Tet) enzymes, a family of enzymes that converts 5-methylcytosine (5mC) to 5-hydroxymethylcytosine (5hmC) [16]. Thus, we used Tet-assisted bisulfite sequencing (TAB-seq) data collected from non-infected DCs [5] to assess if DM sites had significantly different levels of 5hmC as compared to non-DM sites. We found that DM sites (Cluster 3) show high levels of 5hmC even prior to infection (**Figure 1E**; 3.2-fold enrichment compared to non-DM sites; Wilcoxon test; *P* < 1 × 10^−16^), suggesting that DM sites are likely pre-bound by TET enzymes (likely TET2 [17, 18], the most expressed Tet enzyme in DCs (**Supplementary Figure 3**)) and that 5hmC may serve as a stable mark that acts to prime enhancers [19-21].

### Up-regulation of inflammatory genes precedes DNA demethylation

We collected RNA-seq data from matched non-infected and infected samples at each time point, for a total of 34 RNA-seq profiles across time-treatment combinations (mean = 42.2 million reads per sample; **Supplementary Table 1**). The first principal component of the resulting gene expression data accounted for 63% of the variance in our dataset and separated infected and non-infected DCs (**Supplementary Figure 4A**). We found extensive differences in gene expression levels between infected and non-infected DCs: of the 13,956 genes analyzed, 1,987 (14%), 4,371 (31%), 4,591 (33%), and 5,189 (37%) were differentially expressed (DE) at 2, 18, 48, and 72 hours post-infection, respectively (FDR < 0.01 and absolute log2(fold change) > 1; **Supplementary Table 3**). We also collected RNA-seq data in samples stimulated with heat-inactivated MTB and found that, similar to changes in methylation, changes in gene expression in response to live and heat-inactivated MTB were strongly correlated (r ≥ 0.94; **Supplementary Figure 4B**). We next grouped the set of DE genes across the time course (7,457 in total) into 6 distinct temporal expression clusters (**Figure 2A,B**; **Supplementary Table 3**). These clusters cover a variety of differential expression patterns, including genes which show increasing up-regulation over time (DE Cluster 5: Persistent induced; n = 2,091) to genes in which the highest levels of expression occur at 2 or 18 hours followed by a decrease towards basal levels (DE Cluster 4: Early induced (n = 765), and DE Cluster 6: Intermediate induced (n = 839), respectively) (**Figure 2B**). Gene ontology (GO) enrichment analysis revealed that induced genes were strongly enriched among GO terms directly related to immune function, including defense response (FDR = 1.2 × 10^−11^) and response to cytokine (FDR = 8.2 × 10^−12^), whereas repressed genes were primarily enriched for gene sets associated with metabolic processes (**Figure 2C**; **Supplementary Table 4**).

**Figure 2.**
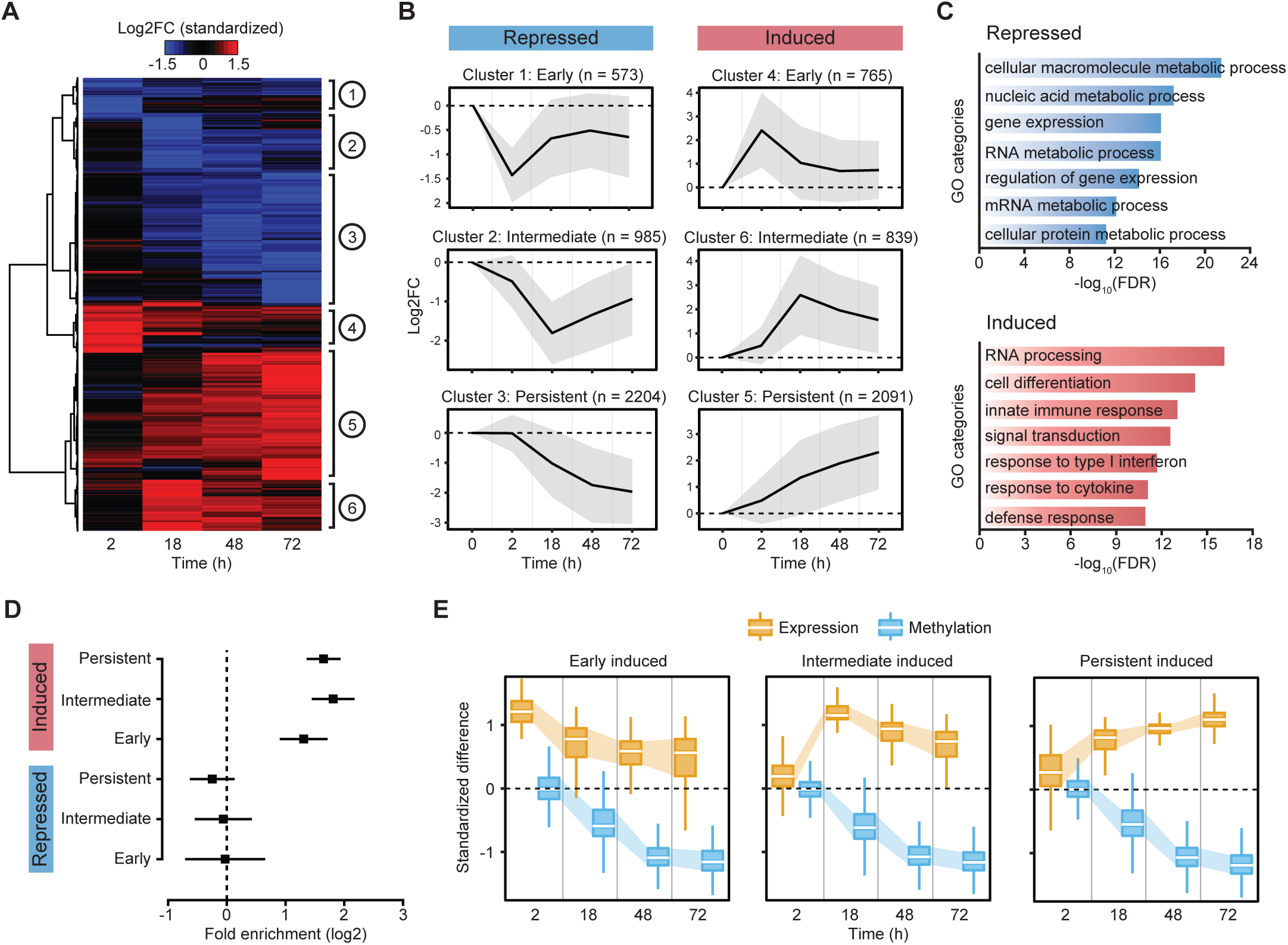
(**A**) Heatmap of differences in expression (standardized log2 fold changes) constructed using unsupervised hierarchical clustering of the 7,457 differentially expressed genes (identified at any time point using cutoffs of |log2FC| > 1 and FDR < 0.01; y-axis) across four time points after MTB infection results in six distinct patterns of changes in expression. (**B**) Mean log2 fold expression changes of genes in each cluster across all time points; shading denotes ±1 standard deviation. For visualization purposes, we also show the ‘0h’ time point, where we expect no changes in expression. (**C**) Gene ontology enrichment analyses among genes that are repressed or induced in response to MTB infection. (**D**) Enrichment (in log2; x-axis) of differentially expressed genes associated with differentially methylated CpG sites (Cluster 3). Error bars show 95% confidence intervals for the enrichment estimates. (**E**) Boxplots showing the distribution of standardized differences in methylation of DM sites in Cluster 3 (blue) along with the corresponding standardized differences in expression of the associated genes (orange), across all time points.

We next tested whether genes located near DM sites—particularly focusing on those sites exhibiting a stable loss of methylation (*i.e.*, Cluster 3 in Figure 1C,D)—were more likely to be differentially expressed upon MTB infection relative to all genes in the genome. We found that genes associated with one or more DM sites were strongly enriched among genes that were up-regulated in response to infection, regardless of the time point at which expression levels started to change: early (2.5-fold, *P* = 3.23 × 10^−11^), intermediate (3.5-fold, *P* = 3.59 × 10^−25^), and persistent (3.1-fold, *P* = 3.80 × 10^−33^) (**Figure 2D,E**).

If demethylation is required for the activation of enhancer elements and the subsequent up-regulation of their target genes, we would expect demethylation to occur *prior* to changes in gene expression; instead, we found the opposite pattern. Among up-regulated genes associated with DM sites (n = 593), 37% exhibited at least a two-fold increase in gene expression levels at 2-hours post-infection, although differential methylation did not begin to be detectable until 18-hours post-infection (**Figure 2E**). For only 17 genes (less than 3% of all up-regulated genes associated with DM sites), DNA demethylation occurred prior to gene activation (**Supplementary Figure 5**), suggesting that no definitive causal relationship between DNA demethylation and gene activation exists.

To confirm that our findings were generalizable to other innate immune cell types and pathogenic infections, we performed a separate time-course analysis of differential methylation in Salmonella-infected macrophages from one additional donor over six time-points (**Supplementary Table 1**). We discovered hundreds of CpG sites that exhibited a progressive loss of methylation over the time course of infection, corroborating our findings in MTB-infected DCs (**Figure 3A**). To assess whether demethylation arises after the activation of associated enhancers, we collected ChIP-seq data for acetylation of histone 3 lysine 27 (H3K27ac) at 2-hours post-infection, as changes in DNA methylation have yet to occur at this point. We found that the deposition of activating H3K27ac marks preceded demethylation at these CpG sites (**Figure 3B**). Moreover, using previously published RNA-seq expression data from Salmonella-infected macrophages [22], we found that most genes associated with these sites were up-regulated at 2-hours post-infection (**Figure 3C**), prior to any changes in methylation. Collectively, these findings indicate that DNA demethylation is not required for the activation of most enhancer elements and that the vast majority of methylation changes induced by infection are a downstream consequence of transcriptional activation.

**Figure 3.**
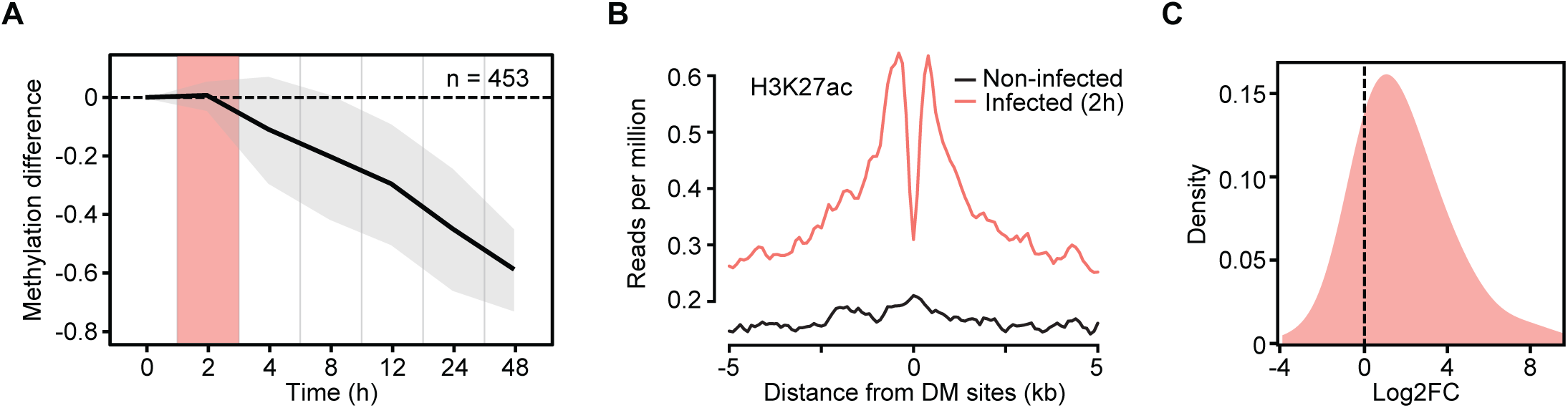
(**A**) Mean differences in methylation (y-axis) in CpG sites that show stable losses of methylation (similar to Cluster 3 DM sites in Figure 1C,D; n = 453) in *Salmonella*-infected macrophages, across six time points after infection (2, 4, 8, 12, 24, and 48 hours (h); x-axis). Shading denotes ±1 standard deviation. For visualization purposes, we also show the ‘0h’ time point, where we expect no changes in methylation. (**B**) Composite plots of patterns of H3K27ac ChIP-seq signals ±5 kb around the midpoints of hypomethylated sites (x-axis) in macrophages at 2 hours post-infection with *Salmonella*. (**C**) Distribution of log2 fold expression changes (between non-infected and *Salmonella*-infected macrophages at 2 hours) for genes associated with DM sites in Figure 3A (n = 269).

### The binding of most infection-induced TFs does not require active demethylation

We next asked whether MTB-induced gene expression changes were associated with changes in chromatin accessibility. To do so, we profiled regions of open chromatin in non-infected and infected DCs at the same time-points (plus one additional time-point at 24 hours) using ATAC-seq [23]. Overall, we found that the response to MTB infection was accompanied by an increase in chromatin accessibility across regulatory regions associated with genes up-regulated upon MTB infection, regardless of their expression profiles (**Figure 4A**). Interestingly, most increases in chromatin accessibility were observed at later stages of infection, suggesting that the activation of early response genes does not require significant modifications to the chromatin structure.

**Figure 4.**
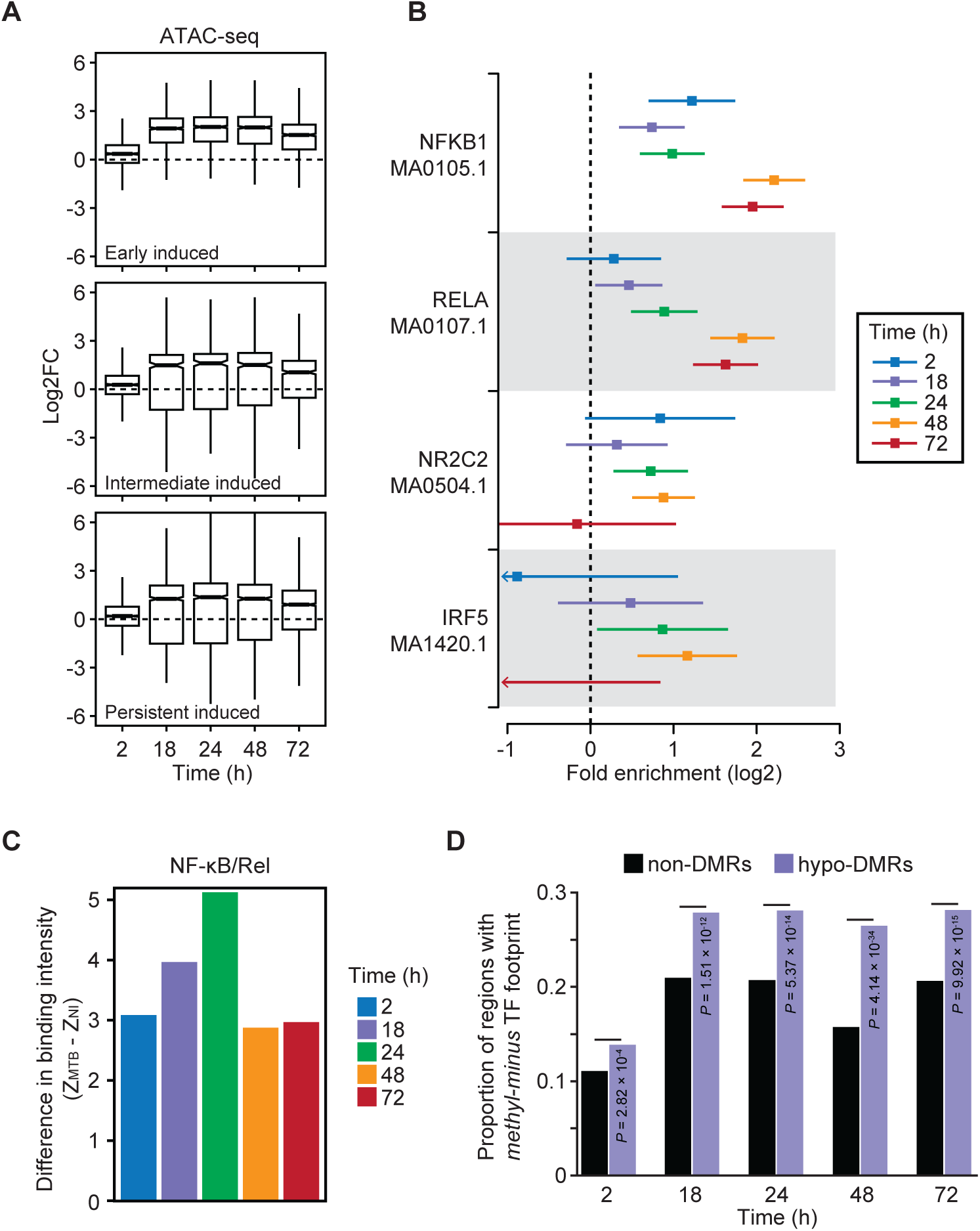
(**A**) Boxplots showing the distribution of log2 fold changes in chromatin accessibility between non-infected and MTB-infected DCs across the five time points of infection (2, 4, 18, 24, 48, and 72 hours) for open chromatin regions associated with the three classes of induced genes described in Figure 2A,B. (**B**) TF binding motifs for which the number of well-supported footprints (posterior probability > 0.99) within hypomethylated regions were enriched (FDR < 0.01) relative to non-DMRs (with 250-bp flanking the start and end) in MTB-infected DCs. The enrichment factors (x-axis) are shown in a log2 scale and error bars reflect the 95% confidence intervals. A complete list of all TF binding motifs for which footprints are enriched within hypomethylated regions can be found in Supplementary Table 5. (**C**) Barplots showing significant differences in TF occupancy score predictions for NF-κB/Rel motifs between MTB-infected and non-infected DCs (Z _MTB_-Z_NI_; y-axis; see **Methods**) across all time points (x-axis). A positive Z-score difference indicates increased TF binding in hypomethylated regions after MTB infection. (**D**) Proportion of regions that overlap a methylation-sensitive (“methyl-minus”; reported in Yin *et al*. [24]) TF footprint (y-axis) observed among non-DMRs and hypomethylated regions (or hypo-DMRs; see **Methods**).

To investigate the relationship between DNA methylation and TF occupancy, we performed TF footprinting analysis on our target regions (*i.e.*, the set of putative enhancers tested for dynamic DNA methylation). We classified target regions as “hypomethylated regions” (n = 1,877) or “non-differentially methylated regions” (non-DMRs) (n = 31,182) according to whether or not these regions overlap DM CpG sites (from differential methylation Cluster 3, specifically). We found that hypomethylated regions were significantly enriched for the binding of immune-related TFs relative to regions exhibiting consistent methylation levels. These immune-related TFs include several master regulators of the innate immune response, such as NF-κB/Rel family members NFKB1 (up to 4.6-fold enrichment across the time course (FDR = 4.78 × 10^−29^)) and RELA (up to 3.6-fold enrichment across the time course (FDR = 1.95 × 10^−18^)) **Figure 4B**; **Supplementary Table 5**).

We next used CentiDual [5] to test for differential binding of TFs between non-infected and infected samples, specifically focusing on the set of TF family members known to orchestrate innate immune responses to infection (*i.e.*, NF-κB/Rel, AP-1, STATs, and IRFs). We found increased binding at NF-κB/Rel binding motifs starting at 2-hours post-infection, despite the fact that no changes in methylation were observed at such early time points (*P* = 0.002; **Figure 4C**; **Supplementary Table 5**; see **Methods**). A similar pattern was observed for AP-1 (*P* = 0.01; **Supplementary Figure 6**). These data show that, while demethylated regions overlap areas bound by immune-induced TFs, the binding of these TFs occurs prior to DNA demethylation.

Although demethylation does not appear to be required for the binding of key TFs involved in regulation of innate immune responses, it is plausible that the removal of methylation marks at DM sites might enable occupancy of methylation-sensitive factors at later time points [24-26]. In support of this hypothesis, we found that, at later time-points (18 hours and above), there was a stronger enrichment for the binding of TFs that preferentially bind to unmethylated motifs (or “methyl-minus” as defined by Yin *et al*. [24]) within hypomethylated regions (up to 1.7-fold enrichment; *χ2*-test; *P* = 4.14 × 10^−34^; **Figure 4D**; see **Methods**). Collectively, these results suggest that, although demethylation is likely not required for the engagement of the core regulatory program induced early after infection, it might play a role in fine-tuning the innate immune response by facilitating the binding of salient methyl-sensitive TFs that mediate later immune responses.

## DISCUSSION

In this study, we generated paired data on DNA methylation, gene expression, and chromatin accessibility in non-infected and MTB-infected DCs at multiple time-points. Our results show that bacterial infection leads to marked remodeling of the methylome of phagocytic cells (both DCs and macrophages), with several thousand CpG sites showing stable losses of methylation via active DNA demethylation.

Strikingly, in our experiment, virtually all changes in gene expression in response to infection occurred prior to detectable alterations in DNA methylation, suggesting that the observed demethylation is a downstream consequence of TF binding and transcriptional activation. We note, however, that our bisulfite sequencing data does not allow us to distinguish between 5mC and 5hmC. Thus, it is possible that the gain of 5hmC in DM sites, which do not show a loss of 5mC at 2-hours post-infection, precedes the activation of certain enhancers, as was recently suggested in T cells [8].

The observed changes in methylation most likely occur via TET2-mediated active demethylation, as previously shown [5, 17, 27]. Consistent with this hypothesis, we found that CpG sites that lose methylation upon infection display high levels of 5hmC at baseline, suggesting that these regions are actively bound by TET2 even prior to infection. Moreover, *TET2* is strongly upregulated 2 hours after infection (∼2.5 fold; **Supplementary Figure 7**). 5hmC could be a stable intermediate that serves as an epigenetic priming mark, ensuring the rapid response of DCs against infection [19-21, 27-30]. Further studies are necessary to investigate the functional relevance of 5hmC in the induction of inflammatory genes during infection.

Using footprint analysis, we show that NF-κB/Rel, a master regulator of inflammation, is recruited to hypomethylated regions as soon as 2-hours post-infection. This finding is consistent with ChIP-seq data collected from macrophages stimulated with Kdo2-Lipid A (KLA), a highly specific TLR4 agonist, which shows that the NF-κB subunit p65 is rapidly recruited to enhancer elements within one hour post-stimulation [31]. We hypothesize that the rapid binding of NF-κB, and of other immune-induced TFs, instigates chromatin opening which is then followed by the recruitment of histone acetyltransferase p300 and the subsequent deposition of activating H3K27ac marks in these regions [32]. Interestingly, p300 can acetylate TET2, conferring enhanced enzyme activity [33], which might account for the eventual loss of DNA methylation in response to infection.

Our results indicate that most changes in gene expression that occur in response to infection are independent of DNA demethylation, further supporting a lack repressive capacity of DNA methylation [34]. Notably, for only 17 genes—out of thousands of differently expressed genes in response to MTB infection—there is evidence that DNA demethylation occurred prior to gene activation. Similar to previous findings [27, 35-40], our results further reinforce the idea that site-specific regulation of DNA demethylation is mediated by TFs that bind to cis-acting sequences. Interestingly, several recent reports have shown that other epigenetic modifications, such as the H3K4me1 enhancer mark, have a similar passive regulatory function [41-43]. However, our results do not exclude the possibility that demethylation might be necessary for the binding of a second wave of TFs that only play a role at later stages of infection (18 hours post-infection or later). In agreement with this hypothesis, we observed a significant enrichment of binding of TFs known to preferentially bind unmethylated CpGs in hypomethylated regions, primarily at later stages post-infection. Ultimately, this suggests that DNA demethylation is not a key regulatory mechanism of early innate immune responses but that it could still play a role in fine-tuning later innate immune responses by facilitating the binding of methylation-sensitive TFs at enhancers.

After an infection is cleared, TFs are expected to unbind, and gene expression as well as DNA methylation levels are anticipated to return to basal state. However, our 72-hour time course study of DNA methylation shows that levels of methylation at DM sites gradually decrease with time post-infection and never revert back to higher levels. Interestingly, this pattern is also observed for genes in which the largest fold changes in gene expression occur at earlier time points. Thus, we speculate that demethylation in response to infection could have a specific biological role in innate immune memory [44-47], and that regions that stably lose methylation may act as primed enhancers, potentially allowing for a faster response to a secondary infection.

## METHODS

### Biological material and sequencing libraries

Buffy coats from healthy donors were purchased from Indiana Blood Center and all participants signed a written consent. The ethics committee at the CHU Sainte-Justine approved the project (protocol #4023). Peripheral blood mononuclear cells (PBMCs) were obtained by centrifugation on Ficoll-Paque, and monocytes were isolated by positive selection with CD14 magnetic beads (Miltenyi Biotec). Monocytes were differentiated into either DCs by adding rhIL-4 (20 ng/mL; Shenandoah Biotechnology,Inc) and rhGM-CSF (20 ng/mL; R&D Systems Inc.) or macrophages by adding rhM-CSF (20ng/mL; R&D Systems Inc.) in the cell culture medium.

DCs were infected with MTB for 2, 18, 48, and 72 h at a multiplicity of infection (MOI) of 1:1 or with heat-killed MTB at MOI of 5:1, as this MOI induces virtually the same transcriptional response at all four time points compared to that observed with live MTB (**Supplementary Figure 2**). Macrophages were infected with *Salmonella typhimurium* as previously described [22]. Briefly, macrophages were infected at MOI of 10:1 for 2 hours, washed, and cultured for 1 hour with 50µg/ml gentamycin, then washed again and cultured in complete medium with 3µg/ml gentamycin for an additional 2, 4, 8, 12, 24 or 48 h, the time points we refer to in the main text.

DNA from DCs was extracted using the PureGene DNA extraction kit (Gentra Systems). DNA from macrophages was extracted using the DNeasy Blood and Tissue Kit (Qiagen). RNA was extracted using the miRNeasy mini kit (Qiagen). RNA quality was evaluated with the 2100 Bioanalyzer (Agilent Technologies) and only samples with no evidence of RNA degradation (RNA integrity number > 8) were kept for further experiments. RNA-seq libraries were prepared using the TruSeq RNA Sample Prep Kit v2, as per the manufacturer’s instructions.

ATAC-seq libraries were generated from 50,000 cells, as previously described [23]. We collected ChIP-seq data for the H3K27ac histone mark in non-infected and *Salmonella*-infected macrophages as previously described [5]. Sequencing was performed using the Illumina HiSeq 2500, as per the manufacturer’s instructions.

### SeqCap Epi library preparation and sequencing

Libraries were generated with KAPA Library Preparation Kit for Illumina Platforms (KAPA Biosystems), as per the manufacturer’s instructions. Briefly, genomic DNA was fragmented to 100-300 bp with an S2 sonicator (Covaris). Fragments were then end-repaired, A-tailed, and ligated with methylated sequencing adapters. Between every enzymatic step, libraries were purified using AMPure beads (Agencourt). After ligation, in addition to the AMPure bead purification, a DUAL-SPRI size selection was performed to further select for fragments with adapters in the window of 200-400 bp. Sodium bisulfite conversion was performed with EZ DNA Methylation Lightning Kit (Zymo Research), and libraries were amplified using KAPA Hifi HotStart Uracil Tolerant Enzyme (KAPA Biosystems). Library quality was assessed by 2100 Bioanalyzer (Agilent Technologies). Samples showing the desired profile were pooled together in equal mass according to Qubit quantification. We then performed a hybridization using the SeqCap Epi kit (Roche NimbleGen). The sample pool, indexes corresponding to the sequences of the adapters used for library preparation, and repetitive DNA (C_0_t) were desiccated and then incubated in hybridization buffer with a set of customized probes for 72 hours to select and sequence target regions only. Specifically, DNA methylation data was collected for 33,059 target regions spanning >130,000 CpG sites (mean length = 300 bp; mean number of CpG sites = 5), which is less than 1% of the ~28 million CpGs contained in the human genome. These regions were primarily comprised of MTB-induced differentially methylated regions identified at 18 hours post-infection using whole-genome bisulfite sequencing, as well as other distal regulatory elements in DCs where changes in DNA methylation have been shown to be most likely to occur (**Supplementary Figure 1A**) [5]. Moreover, these candidate regions were nearby differentially expressed genes in response to MTB at 18 hours. Probes targeting a two kilobase region between coordinates 4500 and 6500 bp of the lambda genome (NC_001416.1) were also included in the SeqCap Epi design, as a control for bisulfite conversion efficiency. Sequencing was performed using the Illumina HiSeq 2500, as per the manufacturer’s instructions.

### SeqCap Epi data processing and differential methylation analysis

Adaptor sequences and low-quality score bases (Phred score < 20) were first trimmed using Trim Galore (http://www.bioinformatics.babraham.ac.uk/projects/trim_galore/). The resulting reads were mapped to the human reference genome (GRCh37/hg19) and lambda phage genome using Bismark [48], which uses Bowtie 2 [49] and a bisulfite converted reference genome for read mapping. Only reads that had a unique alignment were retained. Methylation levels for each CpG site were estimated by counting the number of sequenced C (‘methylated’ reads) divided by the total number of reported C and T (‘unmethylated’ reads) at the same position of the reference genome using Bismark’s methylation extractor tool. We performed a strand-independent analysis of CpG methylation where counts from the two Cs in a CpG and its reverse complement (position *i* on the plus strand and position *i*+1 on the minus strand) were combined and assigned to the position of the C in the plus strand. To assess MethylC-seq bisulfite conversion rate, the frequency of unconverted cytosines (C basecalls) at lambda phage CpG reference positions was calculated from reads uniquely mapped to the lambda phage reference genome. Overall, bisulfite conversion rate was >99% in all of the samples (**Supplementary Table 1**).

In DCs, differentially methylated (DM) CpG sites at each time point following MTB infection were identified using the R package DSS [15]. We used a linear model with the following design: *DNA methylation ~ Donor + Infection*, which allowed us to consider the paired nature of the experiment and capture the effects of infection on DNA methylation observed within donors. We considered a CpG site as differentially methylated if statistically supported at a False Discovery Rate (FDR) < 0.01 and an absolute mean methylation difference above 10%. Only CpG sites that had a coverage of at least 5X in each of the samples were included in the analysis (103,649 in total).

To identify DM sites that show a stable loss of methylation (as Cluster 3 DM sites in DCs) in *Salmonella*-infected macrophages using one individual, we performed a hierarchical clustering analysis on sites that specifically: (*i*) do not change methylation at 2 hours (|methylation difference| < 10%), and (*ii*) lose methylation at 48 hours (methylation difference < −40%).

### 5hmC enrichment at DM sites

To calculate the enrichment of 5-hydroxymethylcytosine (5hmC) at DM CpG sites (Clusters 1, 2 and 3), we compared the distribution of 5hmC levels in non-infected DCs between DM and non-DM sites. Since non-DM sites have lower overall levels of baseline methylation than DM sites **(Supplementary Figure 8A)**, we performed similar enrichment analysis by using a random set of non-DM sites that matches the distribution of methylation found in non-infected samples within each set of DM sites **(Supplementary Figure 8B)**. Each random set contains the same number of CpG sites as those identified within each DM cluster.

### RNA-seq data processing and identification of differentially expressed genes

Read count estimates per gene were obtained using the alignment-free method Kallisto [50]. For all downstream analyses, we excluded non-coding and lowly-expressed genes with an average read count lower than 10 in all of the samples, resulting in 13,955 genes in total. The R package DESeq2 [51] was used to identify differences in expression levels between non-infected and infected samples at each time point. Nominal p-values were corrected for multiple testing using the Benjamini-Hochberg method [52]. The complete list of differentially expressed genes can be found in Supplementary Table 3.

### Gene set enrichment analysis

We used ClueGO [53] at default parameters to test for enrichment of functionally annotated gene sets among differentially expressed genes. The results for these enrichment analyses are reported in Supplementary Table 4. Enrichment p-values were based on a hypergeometric test using the set of 13,955 genes as background. Benjamini-Hochberg method was applied for multiple testing correction.

### ChIP-seq data processing and tag density profiles

ChIP-seq reads were trimmed for adapter sequences and low-quality score bases using Trim Galore. The resulting reads were mapped to the human reference genome using Bowtie 2 with the following option: -N 1. Only reads that had a unique alignment were retained, and PCR duplicates were further removed using Picard tools (http://broadinstitute.github.io/picard/). Tag density profiles for chromatin modifications and genome accessibility patterns around regions of interest were accomplished with ngs.plot package [54] using default parameters.

### ATAC-seq data processing and TF footprinting analysis

ATAC-seq reads were trimmed for adapter sequences and low-quality score bases and were mapped to the human reference genome. Mapping was performed using BWA-MEM [55] in paired-end mode at default parameters. Only reads that had a unique alignment (mapping quality > 10) were retained. TF footprinting analyses were performed as previously described, using the Centidual algorithm [5] and JASPAR annotated human TF binding motifs (2018 release) [56]. For each of the actively bound TFs in DCs (241 in total; FDR < 0.05 at 18 hours post-infection; **Supplementary Table 5**)), we first trained Centidual assuming that the footprint was bound in the two conditions. Then, we fixed the model parameters and generated a likelihood ratio and posterior probability *π_lt_* for each condition *t* separately and for each site *l*. To detect if the footprint was more active in one of the two conditions, we fit a logistic model that included an intercept for each condition (*α* and *δ*), the PWM effect *β*, and PWM times the treatment effect *γ*:

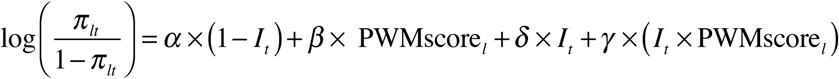

where ***I****_t_* is an indicator variable that takes the value 1 if t = “treatment” and 0 if t = “control”. We then calculated a Z-score for the interaction effect *γ*, corresponding to the evidence for condition-specific binding. ATAC-seq samples were down-sampled to obtain similar number of reads between NI and HI samples at each time-point. We used a window size of 300 bp on either side of the motif match, and reads with fragment lengths [40, 140] and [141, 600] bp for footprinting analyses.

To test for differential binding of immune-related TFs (NF-κB/Rel, AP-1, STATs, and IRFs) between non-infected and infected samples, we compared the intensity of the Tn5 sensitivity-based footprint across all matches to motifs of TFs that belong to each family in the hypomethylated regions. Specifically, the following motif IDs (and corresponding names) were aggregated to their respective TF family:

| NF-κB/Rel |  |
| --- | --- |
| MA0105.3 | NFKB1 |
| MA0105.2 | NFKB1 |
| MA0105.1 | NFKB1 |
| MA0105.4 | NFKB1 |
| MA0778.1 | NFKB2 |
| MA0101.1 | REL |
| MA0107.1 | RELA |
| MA1117.1 | RELB |

| AP-1 |  |
| --- | --- |
| MA0833.1 | ATF4 |
| MA0834.1 | ATF7 |
| MA0462.1 | BATF::JUN |
| MA0835.1 | BATF3 |
| MA0476.1 | FOS |
| MA0099.3 | FOS::JUN |
| MA1126.1 | FOS::JUN(var.2) |
| MA1134.1 | FOS::JUNB |
| MA1141.1 | FOS::JUND |
| MA1127.1 | FOSB::JUN |
| MA1135.1 | FOSB::JUNB |
| MA1136.1 | FOSB::JUNB(var.2) |
| MA0477.1 | FOSL1 |
| MA1128.1 | FOSL1::JUN |
| MA1129.1 | FOSL1::JUN(var.2) |
| MA1137.1 | FOSL1::JUNB |
| MA1142.1 | FOSL1::JUND |
| MA1143.1 | FOSL1::JUND(var.2) |
| MA0478.1 | FOSL2 |
| MA1130.1 | FOSL2::JUN |
| MA1131.1 | FOSL2::JUN(var.2) |
| MA1138.1 | FOSL2::JUNB |
| MA1139.1 | FOSL2::JUNB(var.2) |
| MA1144.1 | FOSL2::JUND |
| MA1145.1 | FOSL2::JUND(var.2) |
| MA0655.1 | JDP2 |
| MA0656.1 | JDP2(var.2) |
| MA0488.1 | JUN |
| MA0489.1 | JUN(var.2) |
| MA1132.1 | JUN::JUNB |
| MA1133.1 | JUN::JUNB(var.2) |
| MA0490.1 | JUNB |
| MA1140.1 | JUNB(var.2) |
| MA0491.1 | JUND |

| STATs |  |
| --- | --- |
| MA0137.1 | STAT1 |
| MA0137.2 | STAT1 |
| MA0137.3 | STAT1 |
| MA0517.1 | STAT1::STAT2 |
| MA0144.2 | STAT3 |

| IRFs |  |
| --- | --- |
| MA0050.2 | IRF1 |
| MA0050.1 | IRF1 |
| MA0051.1 | IRF2 |
| MA1418.1 | IRF3 |
| MA1419.1 | IRF4 |
| MA1420.1 | IRF5 |
| MA0772.1 | IRF7 |
| MA0652.1 | IRF8 |
| MA0653.1 | IRF9 |

To test for enrichment of binding of methylation-sensitive (“methyl-minus”) TFs in hypomethylated regions, we compared the proportion of regions that overlap well-supported footprints (posterior probability > 0.99) of “methyl-minus” TFs reported in Yin *et al*. [24]) among non-DMRs and hypomethylated regions (with 250-bp flanking the start and end). The list of motif IDs (and corresponding names) that were included in the analysis are shown below:

|  |  |  |  |  |  |
| --- | --- | --- | --- | --- | --- |
| MA0018.1 | CREB1 | MA0495.2 | MAFF | MA0823.1 | HEY1 |
| MA0018.3 | CREB1 | MA0511.2 | RUNX2 | MA0830.1 | TCF4 |
| MA0028.2 | ELK1 | MA0526.1 | USF2 | MA0831.1 | TFE3 |
| MA0058.1 | MAX | MA0526.2 | USF2 | MA0831.2 | TFE3 |
| MA0058.3 | MAX | MA0636.1 | BHLHE41 | MA0834.1 | ATF7 |
| MA0059.1 | MAX::MYC | MA0638.1 | CREB3 | MA0835.1 | BATF3 |
| MA0062.1 | GABPA | MA0640.1 | ELF3 | MA0839.1 | CREB3L1 |
| MA0093.2 | USF1 | MA0641.1 | ELF4 | MA0871.1 | TFEC |
| MA0095.1 | YY1 | MA0649.1 | HEY2 | MA1126.1 | FOS::JUN(var.2) |
| MA0095.2 | YY1 | MA0663.1 | MLX | MA1127.1 | FOSB::JUN |
| MA0099.3 | FOS::JUN | MA0664.1 | MLXIPL | MA1128.1 | FOSL1::JUN |
| MA0104.4 | MYCN | MA0736.1 | GLIS2 | MA1129.1 | FOSL1::JUN(var.2) |
| MA0136.2 | ELF5 | MA0749.1 | ZBED1 | MA1130.1 | FOSL2::JUN |
| MA0149.1 | EWSR1-FLI1 | MA0750.1 | ZBTB7A | MA1131.1 | FOSL2::JUN(var.2) |
| MA0156.2 | FEV | MA0750.2 | ZBTB7A | MA1134.1 | FOS::JUNB |
| MA0464.2 | BHLHE40 | MA0757.1 | ONECUT3 | MA1135.1 | FOSB::JUNB |
| MA0470.1 | E2F4 | MA0759.1 | ELK3 | MA1136.1 | FOSB::JUNB(var.2) |
| MA0473.1 | ELF1 | MA0761.1 | ETV1 | MA1137.1 | FOSL1::JUNB |
| MA0473.2 | ELF1 | MA0762.1 | ETV2 | MA1138.1 | FOSL2::JUNB |
| MA0474.2 | ERG | MA0763.1 | ETV3 | MA1139.1 | FOSL2::JUNB(var.2) |
| MA0475.1 | FLI1 | MA0764.1 | ETV4 | MA1141.1 | FOS::JUND |
| MA0475.2 | FLI1 | MA0765.1 | ETV5 | MA1142.1 | FOSL1::JUND |
| MA0476.1 | FOS | MA0772.1 | IRF7 | MA1143.1 | FOSL1::JUND(var.2) |
| MA0477.1 | FOSL1 | MA0821.1 | HES5 | MA1144.1 | FOSL2::JUND |
| MA0478.1 | FOSL2 | MA0822.1 | HES7 | MA1145.1 | FOSL2::JUND(var.2) |

### Relationship between gene expression and chromatin accessibility

Peaks were first called on ATAC-seq using the MACS2 software suite [57] with the added parameters: -g hs -q 0.05 –broad –nomodel – extsize 200 –nolambda. All peaks from each sample were then merged to provide one set of combined peaks. To count the number of reads overlapping peaks, we used featureCount (from the subread package) [58] with the following option: -p -P. For all downstream analyses, we excluded low-count peaks with an average read count lower than 10 across all samples, resulting in 79,282 peaks in total. We then plotted the distribution of changes in Tn5 accessibility (between non-infected and MTB-infected DCs across the five time-points of infection (2, 4, 18, 24, 48, and 72 hours)) for the top 25% most variable peaks associated with DE genes in each cluster. The DE genes associated with the selected peaks represent ~50% of the total genes within each of the DE clusters: (*i*) Early induced: 418/765 = 55%; (*ii*) Intermediate induced: 418/839 = 49%; and (*iii*) Persistent induced: 1083/2091 = 52%.

## DATA ACCESS

Data generated in this study have been submitted to the NCBI Gene Expression Omnibus (GEO; http://www.ncbi.nlm.nih.gov/geo/) under accession numbers **GSE116406** (ATAC-seq), **GSE116411** (ChIP-seq), **GSE116405** (RNA-seq), and **GSE116399** (SeqCap Epi)

## ACKNOWLEDGMENTS

We thank Calcul Quebec and Compute Canada for managing and providing access to the supercomputer Briaree from the University of Montreal. This study was funded by grants from the Canadian Institutes of Health Research (301538 and 232519), and the Canada Research Chairs Program (950-228993) (to L.B.B.). A.P. and F.M-L. were supported by a fellowship from the The Fonds de recherche du Québec – Santé (FRQS).

